# Modality specific memory enhancement in *Heliconius* butterflies

**DOI:** 10.1101/2024.09.14.612954

**Authors:** Elizabeth Hodge, Amaia Alcalde Anton, Louise Bestea, Greta Hernández, Jane Margereth Aguilar, Max S. Farnworth, Denise Dalbasco Dell’Aglio, W. Owen McMillan, Stephen H. Montgomery

**Author notes:** **Correspondence:** EH, AAA, SHM. contributed equally (Elizabeth Hodge, Amaia Alcalde Anton).

## Abstract

How animals perceive, process and respond to environmental cues is tightly tuned to the species-specific demands, and reflected by the structure of neural systems. In the Neotropical butterflies, *Heliconius*, the mushroom bodies, insect learning and memory centres, are significantly expanded compared to their closest relatives. This expansion coincided with the evolution of a novel diet, pollen feeding, and a spatial foraging behaviour consistent with trap-lining. Previous research has shown that *Heliconius* have more accurate visual long-term memory than other Heliconiini. Here, we tested whether this enhanced memory stability is specific to visual cues by conducting a long-term olfactory memory assay in two *Heliconius* species and two outgroup species. We found no differences in the long-term olfactory memory between *Heliconius* species and the outgroup species. Combining data from olfactory and visual memory trials confirms a significant shift in performance among sensory modalities between *Heliconius* and outgroup genera. In contrast, tests of how Heliconiini prioritise olfactory and visual cues when in presented in conflict also show no consistent shift in attentiveness to sensory cues between species. Our data provide a rare case where memory performance has been compared across species and sensory modalities, to identify evidence of a modality specific shift.

## Introduction

How animals perceive, process and respond to the sensory world they inhabit is likely tightly tuned to the species-specific demands imposed by their ecology and life history. Debates concerning the evolution of cognition (broadly defined as the acquisition, processing, storage and retrieval of information) have often focused on the importance of specific or general adaptations – for example through the relative importance of domain general or domain specific cognitive enhancements (Burkart et al., 2017; Poirier et al., 2020), or region-specific adaptations in the brain versus the importance of whole brain size (Barton and Harvey, 2000; Montgomery et al., 2016b). The expansion of specific brain regions can provide specific cognitive advantages (Schumacher and Carlson, 2022; Sukhum et al., 2018; Ward et al., 2012), and some species show adaptive shifts in specific learning and memory tasks (Sherry and Hoshooley, 2010). However, there are few cases where test paradigms have been combined to compare memory performance across species and sensory modalities, to test whether selection shapes the retention and recall of associative memories in a modality specific manner.

In the insect brain, the mushroom bodies are responsible for sensory integration, learning and memory (Menzel and Müller, 1996; Modi et al., 2020; Yu et al., 2006; Zars, 2000). Intrinsic mushroom body neurons, the Kenyon cells, branch out to form the calyx, where they synapse with projection neurons carrying sensory information before extending axonal projections to form the mushroom body lobes. Across insects, differences in species’ sensory ecologies are often reflected in the calyx structure and sensory input (Gronenberg, 2001; Kinoshita et al., 2015; Lin and Strausfeld, 2012; Masuda-Nakagawa et al., 2009), with multisensory input occurring in many species, often within topographically separated calyx regions (Couto et al., 2023; Kinoshita et al., 2015). Changes in connectivity within the mushroom body are thought to be the physical basis of memory formation, and show specific effects on topographical regions of the calyx dependent on the sensory cue (Hourcade et al., 2010). Mushroom body expansion is also observed in several orders, including Hymenoptera, Coleoptera, Dictyoptera and Lepidoptera (Farris, 2013). Within Lepidoptera, the most substantial known expansion of the mushroom bodies occurs in the genus *Heliconius* (Montgomery et al., 2016a). These Neotropical butterflies are part of a larger tribe of butterflies, Heliconiini, found throughout central and south America. Mushroom body size in *Heliconius* is around four times larger than other, closely related outgroup Heliconiini with whom they share habitats and most aspects of their ecology (Couto et al., 2023; Montgomery et al., 2016a; Sivinski, 1989). This volumetric expansion is driven by increased Kenyon cell number, and an increased proportion of calyx that receives visual input (Couto et al., 2023).

Mushroom body expansion in *Heliconius* butterflies has been linked to the evolution of pollen feeding, a novel foraging behaviour not seen in other Lepidopterans (Gilbert, 1972; Young and Montgomery, 2020). *Heliconius* actively collect pollen from a restricted range of plants, providing an adult source of amino acids (Harpel et al., 2015; Krenn et al., 2009). Pollen feeding in *Heliconius* is associated with cognitively demanding foraging behaviours and changes to life-history traits not seen in other genera in Heliconiini (Young and Montgomery, 2020). This includes the evolution of trapline foraging, whereby long-term foraging routes are established between reliable pollen resources (Ehrlich and Gilbert, 1973; Gilbert, 1991; Mallet, 1986). In other insects, trap-line foraging, and allocentric foraging more generally, is supported largely by memorisation of visual landscape cues (Menzel and Greggers, 2015). *Heliconius* are capable spatial learners in both field and insectary conditions (Moura et al., 2023, 2022). Allied to the evolution of pollen feeding, *Heliconius* are also significantly longer lived than other Heliconiini (Gilbert, 1972; Hebberecht et al., 2022), with longer reproductive lifespans (Dunlap-Pianka, 1979; O’Brien et al., 2003), creating the context for increased benefits of long-term memories of foraging routes (Gilbert, 1972). Indeed, they are proficient visual learners (Young and Montgomery, 2023), and possess impressive long-term visual memory (Couto et al., 2023; Young et al., 2024). In previous comparative visual learning assays, all six Heliconiini species tested were able to learn colour associations, but only the *Heliconius* species remembered these associations after eight days, and, in some cases, their memory persisted beyond 13 days (Young et al., 2024). This suggested that *Heliconius’* enhanced visual-memory co-evolved with mushroom body expansion, trap-lining and the evolution of pollen feeding.

*Heliconius*, therefore, present an interesting opportunity for comparative experiments across closely related species with specific anatomical and behavioural differences, but otherwise similar ecologies, to explore the evolution of neural and behavioural change. Here, we compare long-term olfactory memory across four species: two representatives from *Heliconius* (*Heliconius erato demophoon* and *Heliconius melpomene rosina*) and two representatives from outgroup Heliconiini (*Agraulis vanillae* and *Dryas iulia*). Although existing behavioural data strongly suggests that *Heliconius* have superior visual long-term memory in comparison to outgroup Heliconiini (Young et al., 2024), a generalised improvement across sensory modalities has not been ruled out. By mirroring Young et al.’s long-term visual memory experiment (2024), we provide a direct comparison between long-term olfactory and long-term visual memory data to test the hypothesised modality-specific enhancement in *Heliconius’* long-term memory. Using the same two *Heliconius* species (*H. erato* and *H. melpomene)* and *D. iulia* as an outgroup Heliconiini representative, we further test the capacity of Heliconiini to learn visual and olfactory cue combinations. Finally, we then test whether *Heliconius* prioritise visual over olfactory information when making foraging decisions in comparison to the outgroup species, to explore whether variation in memory performance between sensory modalities could reflect general downregulation of attentiveness to one sensory domain.

## Methods

### i. Animal husbandry

Rearing for long term memory assays was performed in January-April 2023. Butterflies of four species within Heliconiini, two *Heliconius* species (*H. erato* and *H. melpomene*) and two outgroup species (*D. iulia* and *A. vanillae*), were raised from stock populations at the Smithsonian Tropical Research Institute insectaries in Gamboa, Panama. Stocks of each species established from local populations were kept in 3×3×2 m mesh cages in ambient conditions and with natural lighting containing pollen resources (*Lantana camara, Palicourea elata* and *Psiguria sp*.) and supplementary artificial feeders containing 20% sugar-water. Larval host plants were introduced several times a week to allow for egg collection. Once hatched, larvae were raised in mesh pop-up cages using the preferred host-plant(s) of each species. *Passiflora biflora* was used for all four species, and *H. erato* and *D. iulia* were raised exclusively on *P. biflora. H. melpomene* were raised on either *P. biflora, Passiflora menispermifolia* or *Passiflora riparia. A. vanillae* were raised on *P. biflora* or *Passiflora tenuifila*. After eclosion, naïve butterflies were moved into the experimental cages. Experimental cages contained artificial feeders and *P. elata* that had had the flowers removed. There were four cages: a pre-training cage (3×3×2 m), two training cages (2×3×2 m), and a test cage (2×3×2 m). Rearing for sensory conflict assays was performed in January-May 2024, in the same manner. These experiments include *D. iulia*, as an outgroup representative, and *H. erato* and *H. melpomene* as representatives from *Heliconius. A. vanillae* stocks were unavailable for this experiment.

### ii. Long-term olfactory memory assay

Long-term olfactory memory was assessed by invoking odour–reward associations using lemongrass (*Cymbopogon schoenanthus* steam-distilled oil; referred to here as lemongrass odour) and orange (*Citrus aurantium dulcis*, cold-pressed oil; referred to here as citrus odour) essential oil solutions (0.66% oil and 99.34% water). We used lemongrass and orange essential oils as Dell’Aglio et al. (2022) found no evidence of a bias towards either the lemongrass or citrus odour in two species of *Heliconius*. Citrus (orange) and lemongrass scents have no known ecological relevance to any of our four species, and are, therefore, presumed to be largely neutral stimuli.

Artificial feeders were made from orange-coloured foam star-shapes that were ∼3 cm in diameter and had a 0.5 ml Eppendorf in the centre that could be filled with sugar–protein solution (2% Critical Care Formula, 20% sugar and 78% water; positive stimulus) or saturated quinine solution (negative stimulus). Feeders were placed in a 4×6 feeder stand, with 12 feeders evenly distributed across each stand. Each artificial feeder was adjacent to at least two 0.5 ml Eppendorf tube odour-wells that were filled with the essential oil solutions, providing a constant odour stimulus across the feeder stand (Figure S1A). Separate feeder stands were used for each odour and were kept at least 1 m apart to minimise odour mixing.

Naïve butterflies were transferred to the pre-training cage on the day of eclosion (between 11:00am and 1:00pm). Each individual was sexed and assigned a unique ID, which was written on both sides of the wing in permanent marker to allow identification. The pre-training cage contained unscented artificial feeders filled with a sugar–amino acid solution. The butterflies were held in this cage for 1–2 full days after eclosion to ensure they were viable and associated the artificial feeders with a food source. After the initial pre-training period (Figure S1A), the butterflies were moved into the test cage for a naïve preference test. The test cage contained two feeder stands, one scented with the citrus odour and the other with the lemongrass odour and empty artificial feeders. Two cameras (GoPro Hero 5) were mounted above each of the feeder stands for two hours between 9am and 11am to record feeding attempts at each stand. The behavioural footage was analysed using BORIS, the Behavioral Observation Research Interactive Software (Friard and Gamba, 2016), to log the number of feeding attempts (defined as a butterfly probing the feeder with the proboscis) and the time at which each attempt was made, for each individual. The naïve preference test was conducted across two days to determine which, if either, of the odours they innately preferred.

After the initial preference test period, butterflies were moved into one of two training cages depending on their behaviour in the initial preference test. In one cage, the citrus odour was positively reinforced (sugar–amino acid solution) and the lemongrass odour negatively reinforced (quinine solution) and, in the other, the lemongrass odour was positively reinforced and citrus odour negatively reinforced. Butterflies were trained on their non-preferred odour, or were randomly assigned to one of the two training cages if they showed no preference in the naïve test. Butterflies remained in the same training cage for four full days. Each day, the reward and non-reward solutions and odour solutions were replaced.

Following the training period, butterflies were moved back to the test cage to establish whether they had learnt the odour–reward association (Figure S1B). The recall test used the same protocol and behavioural analysis as the naïve preference test. After the recall test, the butterflies were moved back to the pre-training cage, where they stayed for eight days with no exposure to the odour stimuli. After an eight-day period, the butterflies were moved a final time to the test cage for the long-term memory (LTM) test, following the same protocol as the other preference tests. An eight-day period was chosen following Young et al.’s (2024) long-term visual memory experiment protocol.

### iii. Conflict assay

Previous protocols have explored how different *Heliconius* species weigh visual and olfactory cues following positive and negative reinforcement (Borrero et al., 2024; Dell’Aglio et al., 2022), based on experimental designs in prior work in hawkmoths (Stöckl et al., 2016). In this experiment, we adapted these protocols to investigate the colour cues used in Young et al. (2024) in combination with the long-term olfactory memory test odour cues above. The experiment used naïve butterflies. After eclosion, butterflies were sexed and ID-ed before being placed in a waiting cage. In the waiting cage, each butterfly was trained to feed from white artificial feeders so that they recognised these as a food source. Butterflies remained in the waiting cage for 1 to 3 days post-eclosion before being transferred to the naïve preference test (Figure S2C). As sensory cues, we used artificial flower feeders made of five-pointed foam stars in two different colours, yellow and purple, with a 0.5 ml Eppendorf tube in the centre, containing either the positive (1% pollen, 20% sugar and 79% water solution) or negative (saturated quinine solution) reinforcement (Figure S2A). Similarly to the long-term olfactory memory assay, the feeders were evenly distributed across a 6×4 feeder stand, which could be introduced and removed from the experimental cages as needed. As an olfactory cue, the feeder stands had evenly distributed odour-wells, which contained one of two essential oil solutions also used in the previous experiment, lemongrass (*Cymbopogon schoenanthus*, steam-distilled oil) and orange (*Citrus aurantium dulcis*, cold-pressed oil), following Dell’Aglio et al. (2022). Each essential oil solution was prepared with 0.66% oil and 99.34% water. This coupled the visual cue of the feeder with a particular odour, such that one combination was positively reinforced, and one was negatively reinforced. As in the long-term memory experiment, the test were recorded using two cameras (GoPro Hero 5) and the videos were annotated using BORIS software (Friard and Gamba, 2016).

After 4-5 days in a waiting cage, butterflies were exposed to two different colour–odour combinations: yellow-lemongrass and purple-citrus. The feeders were left empty, and butterfly behaviour was recorded for two hours to check for individual, naïve preferences. After this test, the butterflies were trained for 3 full days against their initial preference (Figure S2C). They were transferred to a training cage where their initial preference was negatively reinforced, while the non-preferred combination was positively reinforced. If a butterfly did not exhibit a clear preference, one of the training combinations was randomly assigned. After the training period, the butterflies were returned to the test cage, and a recall test was conducted following the same protocol as the naïve preference test. This recall test aimed to determine whether the butterflies had learned the associations during the training period. After the recall test, butterflies were moved back to their training cages for an additional 1 to 2 days to further reinforce the colour–odour associations. On the 14^th^ /15^th^ day, a conflict test was performed, in which the positively reinforced colour was paired with the negatively reinforced odour, and vice versa, so the new combinations were: yellow-citrus and purple-lemongrass (Figure S2B). Following this, butterflies underwent an additional day of the same training they had in the beginning. Finally, a colour preference test was conducted to assess whether the butterflies had learned the odour and colour cues independently (Figure S2C). In this final test, only purple and yellow feeders were presented, with no associated odours.

### iv. Statistical analysis

In the long-term memory assay, as we were interested in retention of the trained association, individuals that did not successfully learn during the training period (accuracy < 50%) were removed from the dataset. This resulted in the removal of 37 individuals: 10 of 65 *A. vanillae*, 13 of 50 *D. iulia*, 8 of 45 *H. erato*, and 6 of 51 *H. melpomene*. Analyses of feeding preference were then carried out using generalised linear mixed models (GLMMs) with a binomial distribution in R v4.3.0 (R Core Team, 2020) using the *glmer* function in the *lme4* package v1.1-33 (Bates et al., 2015). When looking at interspecific differences, species and trial (naïve test, recall test and LTM test) were treated as fixed effects. When looking at intraspecific differences, only trial was used. To determine whether feeding attempts for the two odours were random in the naïve preference test and the LTM test, deviation from 50% was assessed using a null GLMM. The long-term olfactory memory protocol followed Young et al.’s 2024 visual long-term memory experiment as closely as possible. This included using four of the same populations/species to allow comparisons between visual and olfactory long-term memory performance. Although the experiments were performed in different years, the cage structure, environmental conditions, and origin of the butterflies (i.e. from wild derived stocks) were consistent. The data for both experiments were combined and differences in the species’ ability to learn and retain olfactory versus visual associations were analysed using GLMMs. Interspecific differences were analysed using species, trial and experiment (visual or olfactory) as fixed effects and ID and observation-level random effects to account for overdispersion. Additionally, differences in performance between the *Heliconius* genus and outgroup Heliconiini group were analysed, also with trial and experiment as fixed effects.

In the conflict experiment, we aimed to assess the preference for odour or visual cues after the butterflies had learned a colour–odour association. Therefore, we excluded individuals that did not successfully learn this association. During the video analysis, we observed that a few butterflies landed on the odour Eppendorf odour-wells rather than the artificial flowers, seemingly attracted to the source of the odours. We therefore created two datasets: one accounting for these odour-well landings and one that focuses solely on feeding attempts at the artificial flower. The results and analysis from the second dataset are included in the supplementary materials (Figure S4, Table S12, S13). To analyse the conflict data, we used generalised linear mixed models (GLMMs) with a binomial distribution to analyse cue preference, using the *glmer* function from the *lme4* package v1.1-33. To build the model we began with the most complex model and progressively simplified it based on ANOVA results, removing the least significant fixed factor each time. Models were compared using ANOVA, the Akaike Information Criterion (AIC) and Bayesian Information Criterion (BIC). Initially, the model included fixed factors for day and species. We then added the interaction term between day and species, despite its lack of significance, as it was of interest to us. We built additional models to assess whether the proportion of feeding attempts on the trained colour was significantly different from 50% and another model to check for differences in the number of feeding attempts per individual, using a GLMM with a Poisson distribution in this case.

For both experiments, model diagnostics were performed using the *DHARMa* package v0.4.6 and ID and, where necessary, observation-level random effects were included as random effects to account for overdispersion. The package *emmeans* v1.8.6 was used for *post-hoc* pairwise comparisons between species and trial, using the estimated marginal means with Tukey correction for multiple comparisons.

## Results and discussion

### i. Proficient long-term memory of olfactory associations across Heliconiini

*Heliconius* possess expanded mushroom bodies ∼4× larger than their outgroup genera *Dryas* and *Agraulis*. We compared olfactory memory across two *Heliconius* species (*H. erato* and *H. melpomene*) and two outgroup species (*D. iulia* and *A. vanillae*). There was no significant variation between the species in their naive odour preferences (Figure S3, χ^2^ =4.0164, d.f.=3, p = 0.258). *H. melpomene, D. iulia* and *A. vanillae* all showed modestly biased preferences towards the citrus odour (Table S1, S2, Figure S3), but the spread of data covered the full range of feeding preferences in all species. All four species were able to learn the odour associations with high fidelity (Figure S4), showing significant shifts in preference between the naïve test and the recall test (χ^2^ = 400.817, d.f.=1, p <0.001) (Figure 1A, Table S3). There were no differences between the species’ abilities to learn the odour associations (χ^2^ = 3.024, d.f.=3, p= 0.388), with all species performing with similar accuracy. Compared with their performance in the recall test, all species showed a decline in accuracy in the long-term memory test after the eight-day period without exposure to the odour cues (χ^2^ = 57.398, d.f.=1, p < 0.001; Figure 1A, Table S3). This suggests some memory loss, however, all four species showed accuracy greater than chance, suggesting functional long-term olfactory memory lasting over a week (Table S4, Figure 1A). There were no interspecific differences in long-term memory performance (χ^2^ = 2.445, d.f.=3, p = 0.485), and belonging to the *Heliconius* genus did not equate to greater ability to recall the olfactory association (χ^2^ = 0.280, d.f.=1, p = 0.597).

**Figure 1.**
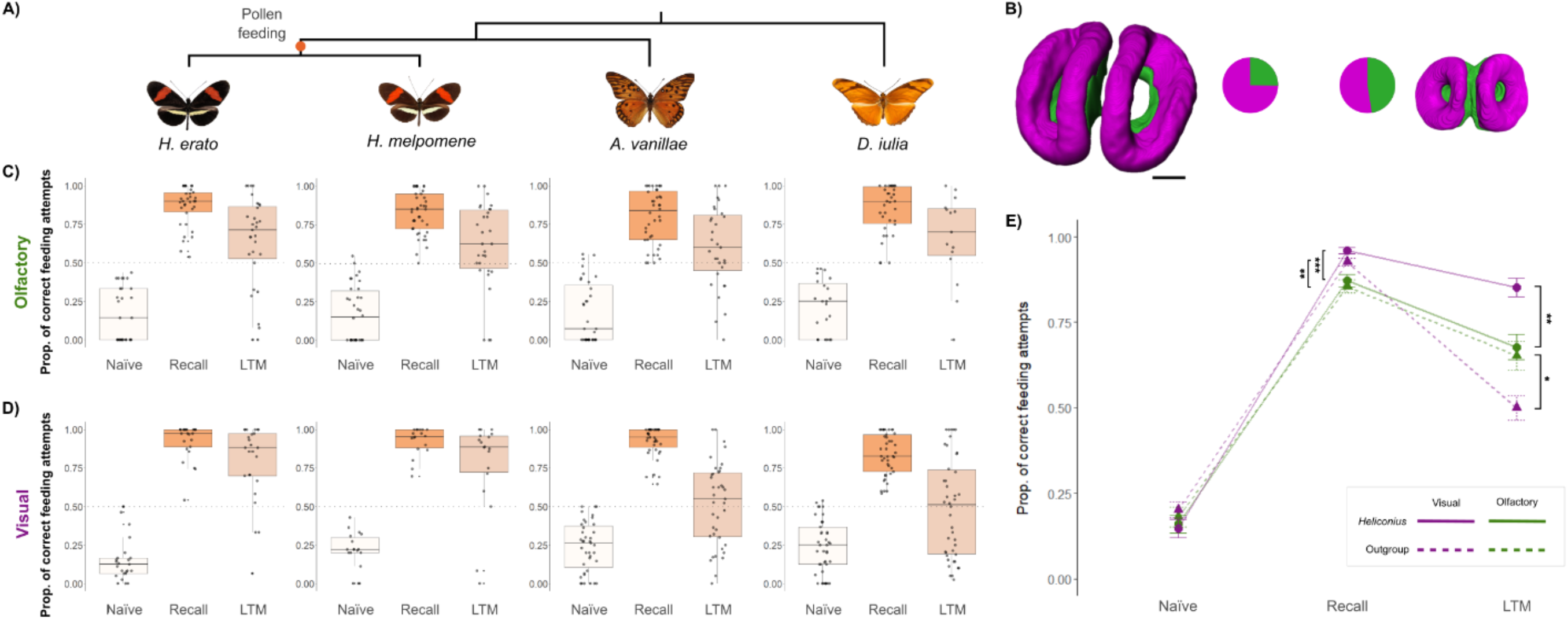
**A**. Phylogeny of species used in the behavioural assays (pollen feeders: *H. erato, H. melpomene*; non-pollen feeders: *A. vanillae* and *D. iulia*). **B**. 3D models of the mushroom body calyx showing the visual processing areas (magenta) and olfactory processing areas (green) in a representative *Heliconius* and outgroup, with their relative proportions shown in pie charts. Scale bar =100μm **C/D**. Performance across long-term olfactory (**C**) and visual (**D**) memory experiments, colour coded by trial: white = naïve preference, dark orange = recall test, light orange = long-term memory (LTM) test after eight days of no reinforcement of the trained cues. **E**. Pooled *Heliconius* species performance (circles with solid line) compared to pooled outgroup Heliconiini species (triangles with dashed line) between the visual (magenta) and olfactory (green) experiments, showing the shift in memory fidelity in *Heliconius*. Values are extracted from a model including a significant interaction between group, *trial* and *experiment*.

Visual cues are thought to be the primary sensory modality exploited during the derived foraging behaviour of *Heliconius*. Nevertheless, olfactory cues are important for all Heliconiini during intraspecific communication, mate choice and detection of close range floral and hostplant cues (Andersson and Dobson, 2003a, 2003b; Estrada et al., 2009; González-Rojas et al., 2020; Kroutov et al., 1999). Our results build on existing evidence of olfactory learning in Heliconiini species: learning of artificial odour cues shown in *H. erato* and *Heliconius himera* (Dell’Aglio et al., 2022), and recognition of vegetative and floral odours shown in *H. melpomene* (Andersson and Dobson, 2003b, 2003a). The mushroom bodies’ role in the formation of olfactory memories and its importance for olfactory processing in insect species is widely recognised (Hourcade et al., 2010, 2009; Menzel and Müller, 1996; Pascual and Préat, 2001). The expansion of *Heliconius* mushroom bodies is primarily driven by an increase in the volume of calyx receiving input from visual projection neurons, with a much lesser expansion of the olfactory calyx. This indicates a degree of visual specialisation in the *Heliconius* mushroom bodies, supported by previous evidence showing increased stability of visual memories in *Heliconius* compared to outgroup Heliconiini (Young et al., 2024). The absence of enhanced olfactory memory may, therefore, suggest a modality specific memory enhancement that mirrors neuroanatomical variation across the tribe.

### ii, Long-term memory recall varies across sensory modalities between pollen feeding and non-pollen feeding Heliconiini

To formally test the hypothesised modality specific shift in memory, we combined data (Table S5) on performance in the olfactory and visual experiments. We found no differences in behaviour between the *Heliconius* and outgroup butterflies in the naïve tests for either experiment (Table S6). In contrast, we found sensory-modality-specific effects on learning and memory. All four species demonstrated comparable or more accurate visual learning than olfactory learning (χ^2^ = 28.331, d.f.=1, p < 0.001; Table S6, S7). *Heliconius*’ long-term visual memory was also superior to their olfactory long-term memory (Table S6; Figure 1C), and to the long-term visual memory of the outgroup genera (as reported in Young et al., 2024; Table S6). In contrast, the outgroup genera had superior olfactory memory to visual memory (Table S6), and no group differences were found in olfactory memory (χ^2^ = 2.579, d.f.=1, p = 0.612; Figure 1C). These group level effects are borne out in patterns of intra-specific performance (Table S7, Figure 1A,B). Both *Heliconius* species were more accurate in the visual recall test than in the olfactory recall test, and had more accurate long-term visual memory than long-term olfactory memory (Table S6, S7, Figure S5). In contrast, there were no differences in accuracy in learning olfactory versus visual cues for *D. iulia* (Table S7), which showed higher accuracy in the olfactory long-term memory test (Figure S5). In the recall test, *A* .*vanillae* were better at learning visual associations than olfactory associations, but performed equally well in both long-term memory tests (Table S7). While the intensive nature of performing these experiments across multiple, free-flying species restricts the range of stimuli used, we focused on ecologically neutral odours, and equally non-preferred colours. Given learning performance in the recall test are largely consistent across species, and the salience of each association with the reward is equal, we argue the learned associations are typical of general performance.

### iii. No consistent shifts in prioritisation of visual and olfactory cues during foraging decisions by Heliconiini

We next tested how Heliconiini butterflies prioritise visual or olfactory cues when presented in conflict to test whether the apparent shift in visual memory stability was specific to learnt preferences or reflected changes in sensory attention between *Heliconius* and other Heliconiini. All three species successfully learnt the colour–odour association when trained against their initial preference (see recall test in Figure 2; Table S8, S9). The cues were subsequently presented in conflict, with the positively reinforced colour now paired with the negatively reinforced odour. With these conflicting cues, there were significant differences between the proportion of feeding attempts made towards the positively reinforced colour between the recall test and the conflict test (Table S8). This suggests both visual and olfactory cues are important when making foraging decisions in Heliconiini. We also observed an increase in the variability of responses, which likely reflects a greater degree of ‘uncertainty’ during decision making. In the conflict test, the proportion of feeding attempts on the positively reinforced colour was significantly above 50% in *H. erato*, suggesting a preference for visual cues over olfactory ones (Table S10). A similar trend was observed in *H. melpomene* butterflies, though this was not significant (Table S10, Figure 2). In the case of *D. iulia*, there appeared to be no preference for either cue (Table S10). However, overall, we found no significant differences between species during the conflict test (Table S9). Significant differences were observed between the conflict test and the colour test for *Heliconius* butterflies, but not for *D. iulia* (*erato*: estimate=-1.334, z.ratio=-2.435, p=0.040; *melpomene*: estimate=-2.157, z.ratio=-2.440, p=0.039; and *Dryas*: estimate=-0.766, z.ratio=-0.922, p=0.626; Table S8). Finally, *Heliconius* also tended to perform better during the final colour test, where the odour cues were absent, but this difference was not significant (Table S9, Figure 2). We also note that *D. iulia* individuals, made significantly fewer feeding attempts per individual in the later trials, particularly during the conflict test (*D*.*iulia* vs. *H. erato*: estimate = -1.28, z.ratio = -5.627, p<0.001 and *D. iulia* vs. *H. Melpomene*: estimate = -1.211, z.ratio = -4.332, p<0.001; Table S11), which may indicate greater uncertainty in decision making. Overall, these results suggest that both visual and olfactory cues are extracted by Heliconiini butterflies, and used with similar importance across genera. While there is some suggestion *Heliconius* may rely more on visual cues when learning colour–odour combinations, this was not sufficient to drive differences in performance when the cues were presented in conflict.

**Figure 2.**
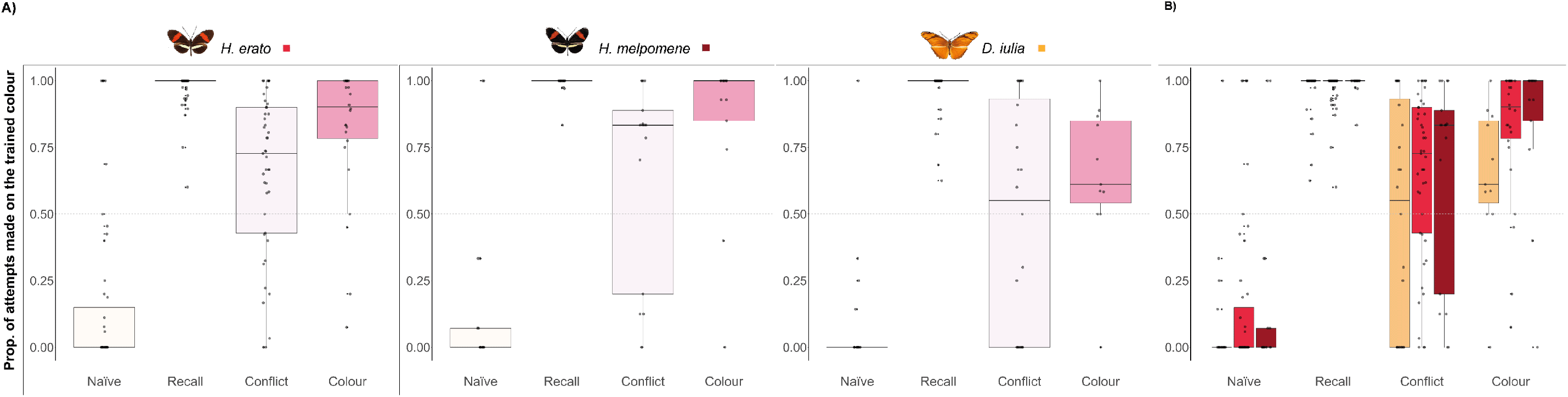
Proportion of feeding attempts made on the trained colour by *H. erato, H. melpomene*, and *D. iulia* across the different tests. The graphs are colour-coded by test: white for naïve preference, dark orange for the recall test, light pink for the conflict test, and pink for the colour-only test. In the last graph is colour-coded by species, orange for *D. iulia*, light red for *H. erato* and red for *H. melpomene*. (**A**) Shows the performance of each species across the different tests. (**B**) Compares the performance of the three species across the four days.

### iv. Visual specialisation of memory performance and circuitry in Heliconius

Our results support the hypothesis that increased investment in visual processing in the mushroom body calyx is linked to specific, derived behaviours of *Heliconius*. It is likely that, when trap-lining, *Heliconius* use a combination of sensory stimuli but rely on visual cues for navigation and continued daily use of foraging routes (Moura et al., 2023; Moura et al., in review). As these routes can be utilised for many weeks or months (Gilbert, 2014; Mallet, 1986; Young and Montgomery, 2020), *Heliconius* may have experienced increased selection for greater retention of visual long-term memories. Indeed, our data suggest a modality specific memory enhancement, specifically increased long-term stability of visual memories over olfactory memories. Our conflict experiments further suggest that this occurs in the absence of a consistent shift in how visual and olfactory associations are prioritised during memory formation, implying that the greater retention of visual memories in *Heliconius* is unlikely due to any shift in sensory reception.

Our conclusions align with both the inferred conservation of sensory pathways across pollen feeding and non-pollen feeding Heliconiini (Hodge et al., in prep.), and with the specific expansion of visual processing regions of the mushroom body calyx, which dominate the volumetric increase observed in *Heliconius* (Couto et al., 2023). *Heliconius* have a 6–8-fold increase in Kenyon cell number and, whilst the proportion of cells receiving visual versus olfactory input has not been clearly delineated, it is likely that these additional cells predominantly receive visual information. Evidence also suggests that a change in the degree of post-eclosion synaptic plasticity accompanies this Kenyon cell increase. *Heliconius* have a greater degree of, and environmental sensitivity to, synaptic pruning compared to *D. iulia* (Young et al., 2024). In Hymenoptera, synaptic plasticity in the calyx is linked to long-term memory formation (Hourcade et al., 2010) and occurs in a sensory modality specific manner, with olfactory experience predominantly effecting the Hymenopteran calyx lip, while visual experience affects the collar (Gronenberg, 2001; Hourcade et al., 2010). To our knowledge, no study in insects has reported evidence of an evolutionary changes in memory performance that is specific to a sensory modality. Whether the improved visual long-term memory in *Heliconius* is directly linked to increased Kenyon cell number, or plasticity, remains to be determined. However, the absence of detectable changes in upstream sensory pathways (Couto et al., 2023) strongly implies these behavioural changes are based in altered mushroom body function.

### iv. Summary

To our knowledge, we have demonstrated the first case of an evolved, memory enhancement that is specific to certain sensory modalities in any insect. This specificity likely reflects the lack of modality general mechanisms for increasing memory enhancement, perhaps due to the high cost of long-term memory (Mery and Kawecki, 2005). Given that long-range navigation of learned spatial routes is supported by visual cues in insects (Collett and Collett, 2002; Graham and Philippides, 2017), and the likely greater stability of landscape cues compared to olfactory plumes in tropical forests, we suggest the enhanced long-term memory of visual cues is adaptive for *Heliconius* as it improves foraging efficiency, while similar changes in olfactory memory would be less beneficial. Combined with neuroanatomical evidence for visual specialisation of the mushroom body, our results provide evidence that selection shapes specific aspects of cognitive performance to meet a species’ ecological learning and memory needs.

## Acknowledgements

We thank the Ministerio del Ambiente, Panama, and the science support, administrative, and facilities staff at the Smithsonian Tropical Research Institute in Panama for making this research possible. In particular, we thank Rémi Mauxion, Oscar Paneso, Cruz Batista and Laura Hebberecht for help and advice in butterfly stock and host plant care, and Fletcher Young and to Jessica Foley for helpful discussions on experimental design and analyses. This work was supported by a NERC Independent Research Fellowship (NE/N014936/1) and an ERC Starter Grant (758508) to SHM and the Smithsonian Tropical Research Institute.

