## Supplementary Information for "Modality specific memory enhancement in *Heliconius* butterflies"

**Institutions:**

[1] University of Bristol, School of Biological Sciences, Bristol, UK, BS8 1TQ

[2] Smithsonian Tropical Research Institute, Gamboa, Panama

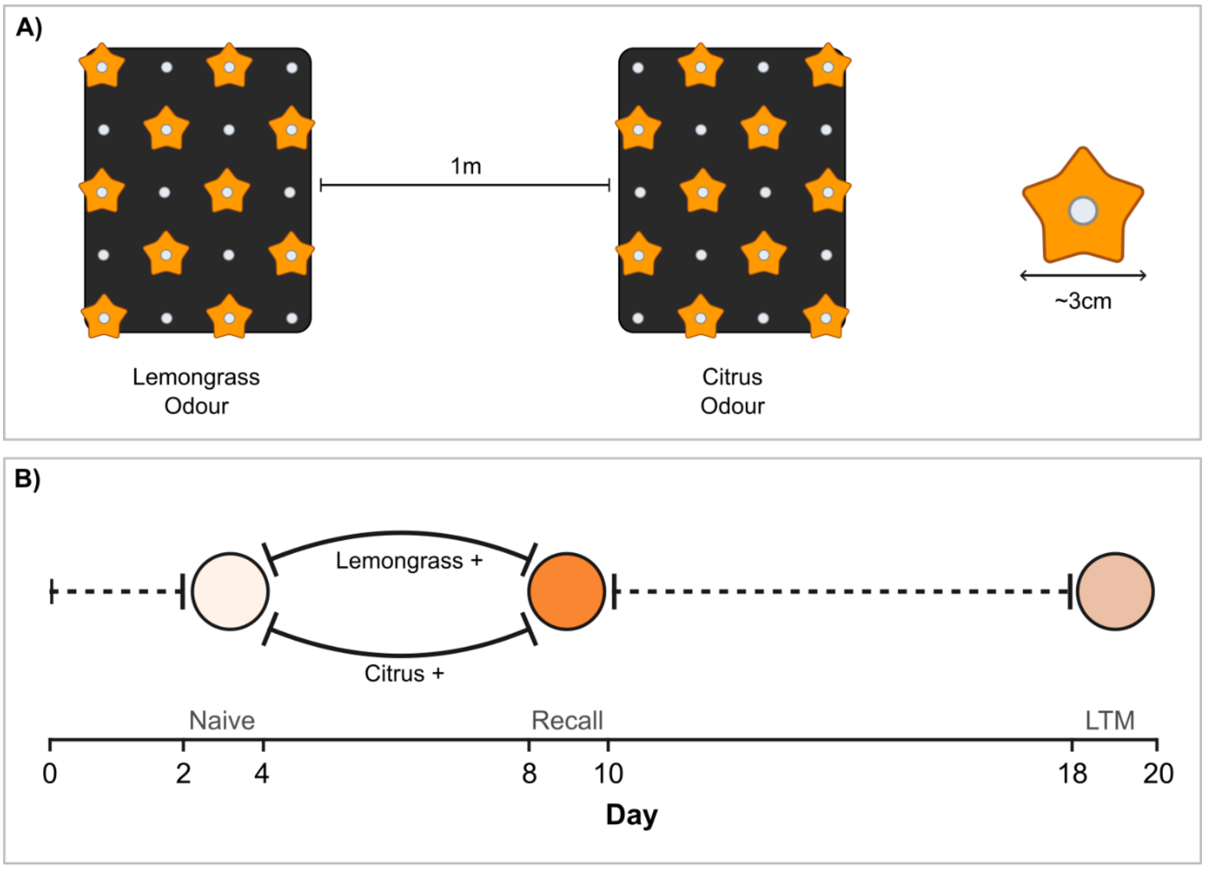


**Figure S1.** Experimental setup (**A**) and protocol timeline (**B**) for the olfactory long-term memory experiment. (**A**) Feeder stand set-up: During the training, the artificial feeders (orange stars) are reinforced with sugar and quinine solutions. During the preference tests, the feeders are empty. Odour wells (white dots) between the two feeders are filled exclusively with either citrus odour or lemongrass odour solutions. (**B**) The bold line indicates the training period, and dashed lines indicate waiting periods where odours are not reinforced. Circles indicate the recorded preference tests.

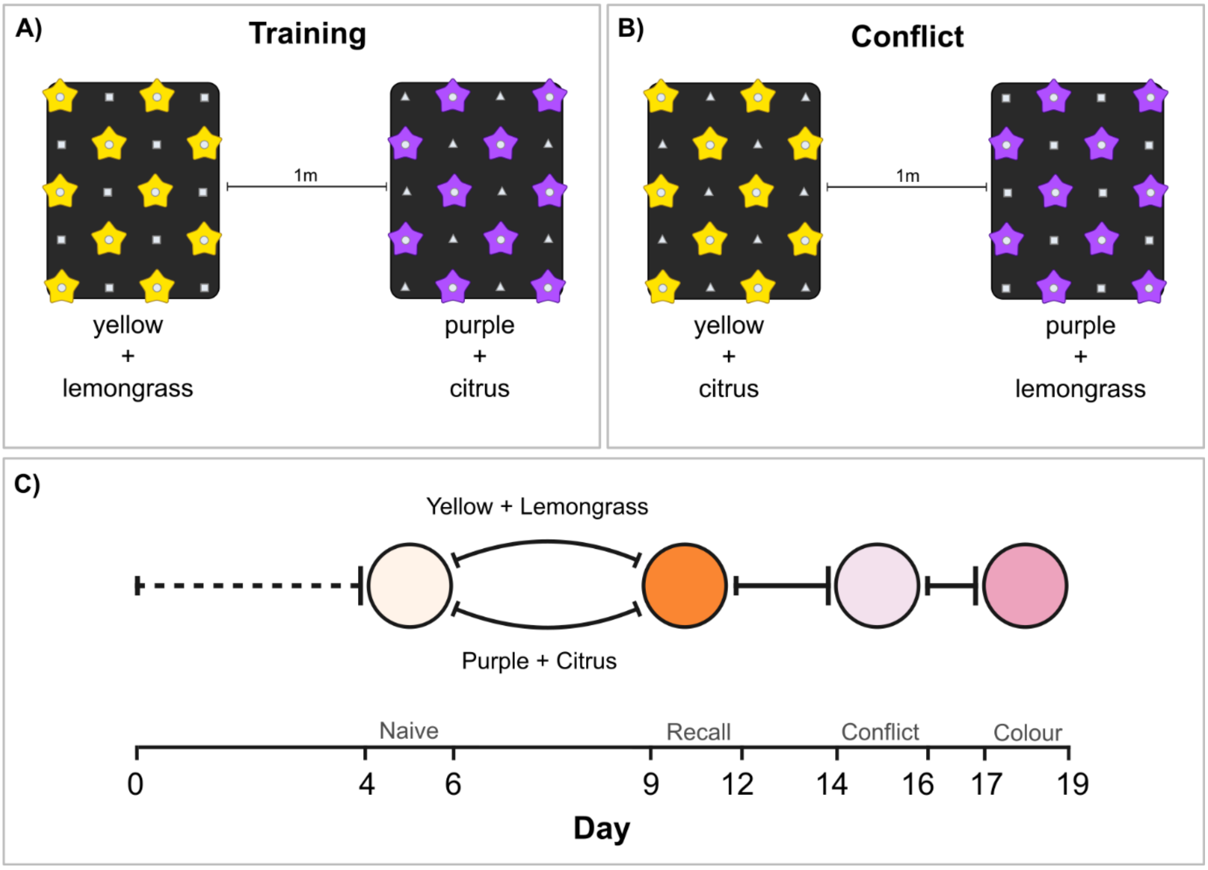


**Figure S2.** Experimental setup for the training (**A**) and conflict test (**B**), and the timeline for the full conflict experiment (**C**). (**A**) Feeder stand set-up for the training: the artificial feeders (yellow or purple stars, depending on the naive preference test) are reinforced with sugar and quinine solutions. The odour wells are filled with either lemongrass odour (squares) or citrus odour (triangles) solutions. (**B**) Feeder stand set-up for the conflict test: the artificial feeders are empty during the preference tests, and the positively reinforced colour is paired with the negatively reinforced odour and vice versa. (**C**) Bold lines indicate the training periods, and dashed lines indicate waiting periods where odours are not reinforced. Circles indicate the recorded preference tests.

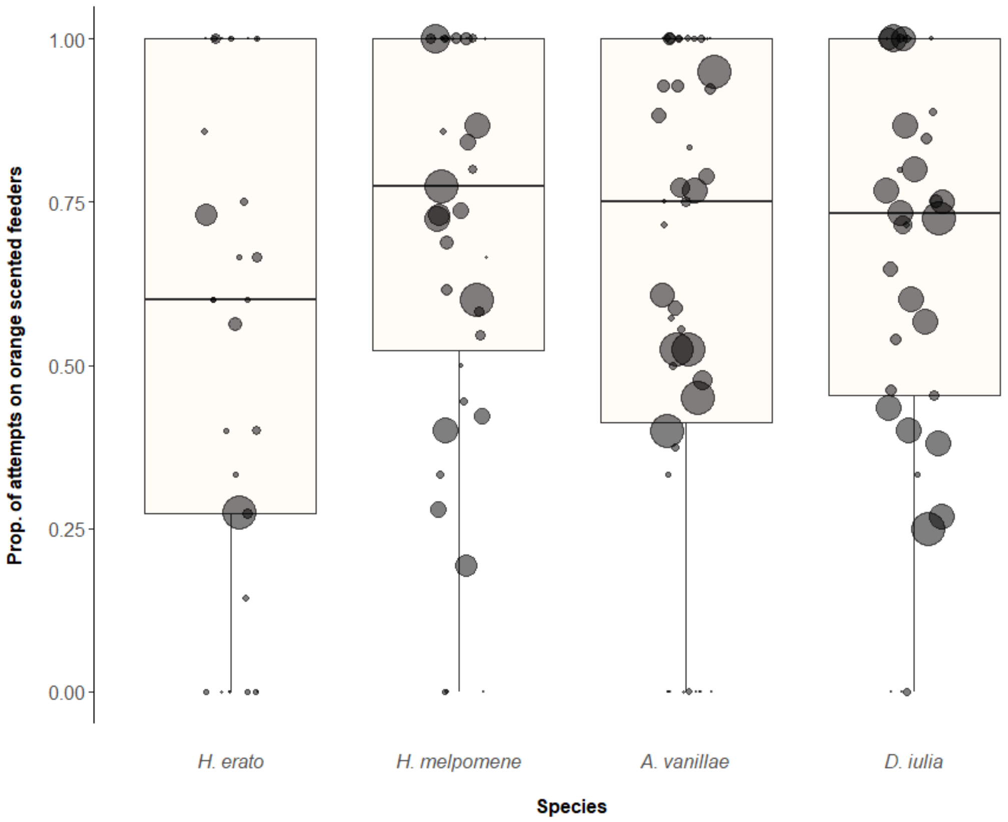


**Figure S3.** Naïve preference for the experimental odours, where 1 equates to total preference for citrus (orange) odour and zero equates to total preference for lemongrass odour. *H. melpomene* (n = 39), *H. erato* (n = 33), *A. vanillae* (n = 46) and *D. iulia* (n = 37). Point size is scaled with number of feeding attempts made by an individual.

**Table S1.** Comparison to 50% in the proportion of feeding attempts on each odour in the naïve preference test for the experimental odours in the long-term olfactory memory experiment. Calculated using a null GLMM only involving ID as a random effect. Species with preference different to 50% are indicated by $p < 0.05$.

| Species | z value | Pr(>\|z\|) |
|---|---|---|
| *H. erato* | -0.929 | 0.353 |
| *H. melpomene* | -3.631 | **<0.001** |
| *D. iulia* | -3.400 | **<0.001** |
| *A. vanillae* | -3.686 | **<0.001** |

**Table S2.** Pairwise comparisons with Tukey correction of the impact of the odour used in the training on the recall performance when learners with accuracy lower than 50% are removed.

| Species | Contrast | z. ratio | p. value |
|---|---|---|---|
| *H. erato* | Lemon - Orange | -0.978 | 0.328 |
| *H. melpomene* | Lemon - Orange | 0.189 | 0.850 |
| *D. iulia* | Lemon - Orange | 0.159 | 0.874 |
| *A. vanillae* | Lemon - Orange | -3.393 | **< 0.001** |

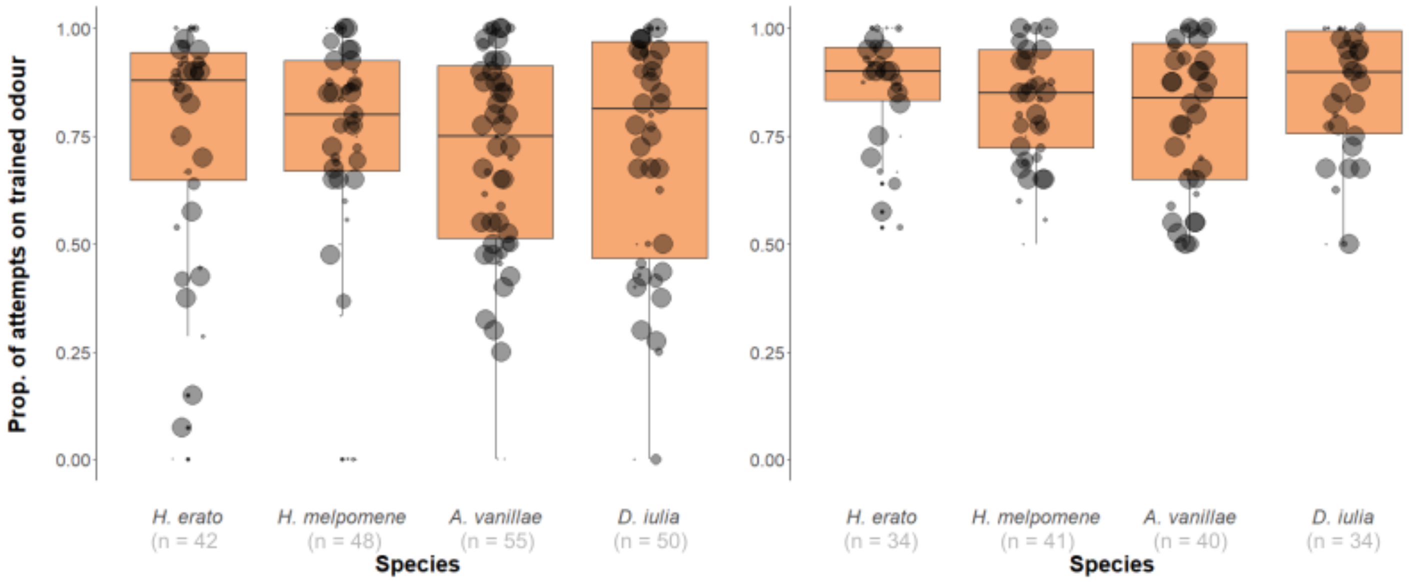


**Figure S4.** Proportion of feeding attempts made by *H. erato, H. melpomene, A. vanillae* and *D. iulia* on the trained odour during the recall test in the long-term olfactory memory test. (Left) Recall performance of all individuals before filtering out unsuccessful learners. (Right) As left panel but with individuals that did not learn the association (accuracy below 50%) removed. There were no significant differences in learning performance before data filtering ($\chi^2$ = 1.524, d.f.=3, p= 0.679) or after filtering ($\chi^2$ = 3.024, d.f.=3, p= 0.388). When learners of accuracy below 60% and 70% were removed, training still did not significantly explain variation in the data (60%: $\chi^2$ = 2.314, d.f.= 1, p = 0.128; 70%: $\chi^2$ = 3.667 d.f.= 1, p = 0.056) (see also Table S4, S5). Point size is scaled with number of feeding attempts made by an individual.

**Table S3.** Pairwise comparisons with Tukey correction of intraspecific shifts in odour preference between the different stages of the long-term memory test. Percentages indicate level at which individuals that did not learn were filtered out (50%, 60% and 70%).

| | | 50% | | 60% | | 70% | |
|---|---|---|---|---|---|---|---|
| **Species** | **Contrast** | **z. ratio** | **p. value** | **z. ratio** | **p. value** | **z. ratio** | **p. value** |
| *H. erato* | Naïve – Recall | -15.085 | **< 0.0001** | -14.999 | **< 0.0001** | -14.997 | **< 0.0001** |
| | LTM – Recall | 7.045 | **< 0.0001** | -6.637 | **< 0.0001** | -6.595 | **< 0.0001** |
| *H. melpomene* | Naïve – Recall | -18.292 | **< 0.0001** | -18.267 | **< 0.0001** | -17.523 | **< 0.0001** |
| | LTM – Recall | 6.129 | **< 0.0001** | -6.004 | **< 0.0001** | -7.556 | **< 0.0001** |
| *D. iulia* | Naïve – Recall | -16.534 | **< 0.0001** | -16.515 | **< 0.0001** | -16.223 | **< 0.0001** |
| | LTM – Recall | 4.892 | **< 0.0001** | -5.372 | **< 0.0001** | -7.126 | **< 0.0001** |
| *A. vanillae* | Naïve – Recall | -10.301 | **< 0.0001** | -17.671 | **< 0.0001** | -16.678 | **< 0.0001** |
| | LTM – Recall | 4.253 | **< 0.0001** | -8.642 | **< 0.0001** | -10.272 | **< 0.0001** |

**Table S4.** Comparison to 50% in the proportion of correct feeding attempts on the trained odour in the long-term memory test in the olfactory memory experiment. Calculated using a null GLMM only involving ID as a random effect. Species with a difference in accuracy to 50% are indicated by $p < 0.05$. Percentages indicate level at which individuals that did not learn were filtered out (50%, 60% and 70%).

| | 50% | | 60% | | 70% | |
|---|---|---|---|---|---|---|
| **Species** | **z value** | **Pr(>\|z\|)** | **z value** | **Pr(>\|z\|)** | **z value** | **Pr(>\|z\|)** |
| *H. erato* | 3.003 | **< 0.01** | 3.567 | **< 0.001** | 4.399 | **< 0.0001** |
| *H. melpomene* | 4.196 | **< 0.0001** | 4.274 | **< 0.0001** | 3.488 | **< 0.001** |
| *D. iulia* | 3.174 | **< 0.01** | 2.796 | **< 0.01** | 2.715 | **< 0.01** |
| *A. vanillae* | 2.234 | **< 0.05** | 2.859 | **< 0.01** | 2.352 | **< 0.05** |

**Table S5.** Number of individuals in each stage of the experiments. Each individual completed all three test stages. N numbers indicate the number of individuals that made feeding attempts in each stage, with individuals that performed with less than 50%, 60% and 70% accuracy in the recall test filtered out.

| | | | n | | |
|---|---|---|---|---|---|
| **Species** | **Experiment** | **Trial** | **50%** | **60%** | **70%** |
| ***A. vanillae*** | Olfactory | Naïve | 35 | 28 | 23 |
| | | Recall | 40 | 32 | 27 |
| | | LTM | 31 | 26 | 24 |
| | Visual | Naïve | 39 | 39 | 36 |
| | | Recall | 39 | 39 | 36 |
| | | LTM | 39 | 39 | 36 |
| ***D. iulia*** | Olfactory | Naïve | 23 | 22 | 19 |
| | | Recall | 34 | 31 | 27 |
| | | LTM | 15 | 14 | 13 |
| | Visual | Naïve | 41 | 40 | 34 |
| | | Recall | 41 | 40 | 34 |
| | | LTM | 42 | 41 | 35 |
| ***H. erato*** | Olfactory | Naïve | 29 | 27 | 25 |
| | | Recall | 34 | 32 | 29 |
| | | LTM | 31 | 29 | 28 |
| | Visual | Naïve | 25 | 24 | 24 |
| | | Recall | 25 | 24 | 24 |
| | | LTM | 25 | 24 | 24 |
| ***H. melpomene*** | Olfactory | Naïve | 34 | 32 | 27 |
| | | Recall | 41 | 39 | 32 |
| | | LTM | 30 | 28 | 23 |
| | Visual | Naïve | 18 | 18 | 17 |
| | | Recall | 18 | 18 | 17 |
| | | LTM | 18 | 18 | 17 |

**Table S6.** Within group (*Heliconius* and outgroup) differences in performance between the olfactory and visual experiments in the different test stages. (N: outgroup; Y: *Heliconius*). Percentages indicate level at which individuals that did not learn were filtered out (50%, 60% and 70%).

| | | **50%** | | **60%** | | **70%** | |
|---|---|---|---|---|---|---|---|
| **Trial** | **Contrast** | **z. ratio** | **p. value** | **z. ratio** | **p. value** | **z. ratio** | **p. value** |
| Naïve | N Olfactory – N Visual | -0.533 | 0.996 | -0.043 | 1.0 | -0.146 | 1.000 |
| | N Olfactory – Y Visual | 0.869 | 0.946 | 1.246 | 0.762 | 1.308 | 0.720 |
| | Y Olfactory – N Visual | -1.154 | 0.820 | -1.230 | 0.773 | -1.742 | 0.399 |
| | Y Olfactory – Y visual | 0.323 | 0.999 | 0.233 | 1.0 | -0.104 | 1.000 |
| Recall | N Olfactory – N Visual | -3.422 | **< 0.01** | -1.927 | 0.283 | -1.724 | 0.412 |
| | N Olfactory – Y Visual | -4.922 | **< 0.0001** | -3.970 | **< 0.001** | -3.038 | **< 0.05** |
| | Y Olfactory – N Visual | -2.707 | **< 0.05** | -2.406 | 0.093 | -2.361 | 0.105 |
| | Y Olfactory – Y visual | -4.353 | **< 0.0001** | -4.361 | **< 0.0001** | -3.569 | **< 0.01** |
| LTM | N Olfactory – N Visual | 2.618 | **< 0.05** | 3.091 | **< 0.05** | 2.167 | 0.168 |
| | N Olfactory – Y Visual | -3.981 | **< 0.001** | -3.386 | **< 0.01** | -4.156 | **< 0.001** |
| | Y Olfactory – N Visual | 3.356 | **< 0.01** | 3.377 | **< 0.001** | 3.388 | **< 0.01** |
| | Y Olfactory – Y visual | -3.696 | **< 0.01** | -3.250 | **< 0.01** | -3.442 | **< 0.01** |

**Table S7.** Interspecific differences in performance between the visual experiment and olfactory experiment in the recall and long-term memory tests. Percentages indicate level at which individuals that did not learn were filtered out (50%, 60% and 70%).

| | | | **50%** | | **60%** | | **70%** | |
|---|---|---|---|---|---|---|---|---|
| **Species** | **Contrast** | **Trial** | **z. ratio** | **p. value** | **z. ratio** | **p. value** | **z. ratio** | **p. value** |
| ***H. erato*** | Olfactory – Visual | Recall | -2.951 | **< 0.01** | -3.055 | **< 0.01** | -2.601 | **< 0.01** |
| | | LTM | -2.498 | **< 0.05** | -2.023 | **< 0.05** | -1.697 | 0.090 |
| ***H. melpomene*** | Olfactory – Visual | Recall | -2.641 | **< 0.01** | -2.474 | **< 0.05** | -1.953 | 0.051 |
| | | LTM | -2.447 | **< 0.05** | -2.340 | **< 0.05** | -3.059 | **< 0.01** |
| ***D. iulia*** | Olfactory – Visual | Recall | -0.487 | 0.626 | -0.039 | 0.969 | -0.084 | 0.933 |
| | | LTM | 2.622 | **< 0.01** | 2.526 | **< 0.05** | 1.729 | 0.084 |
| ***A. vanillae*** | Olfactory – Visual | Recall | -4.201 | **< 0.0001** | -2.644 | **< 0.01** | -2.211 | **< 0.05** |
| | | LTM | 1.083 | 0.279 | 1.556 | 0.120 | 1.134 | 0.257 |

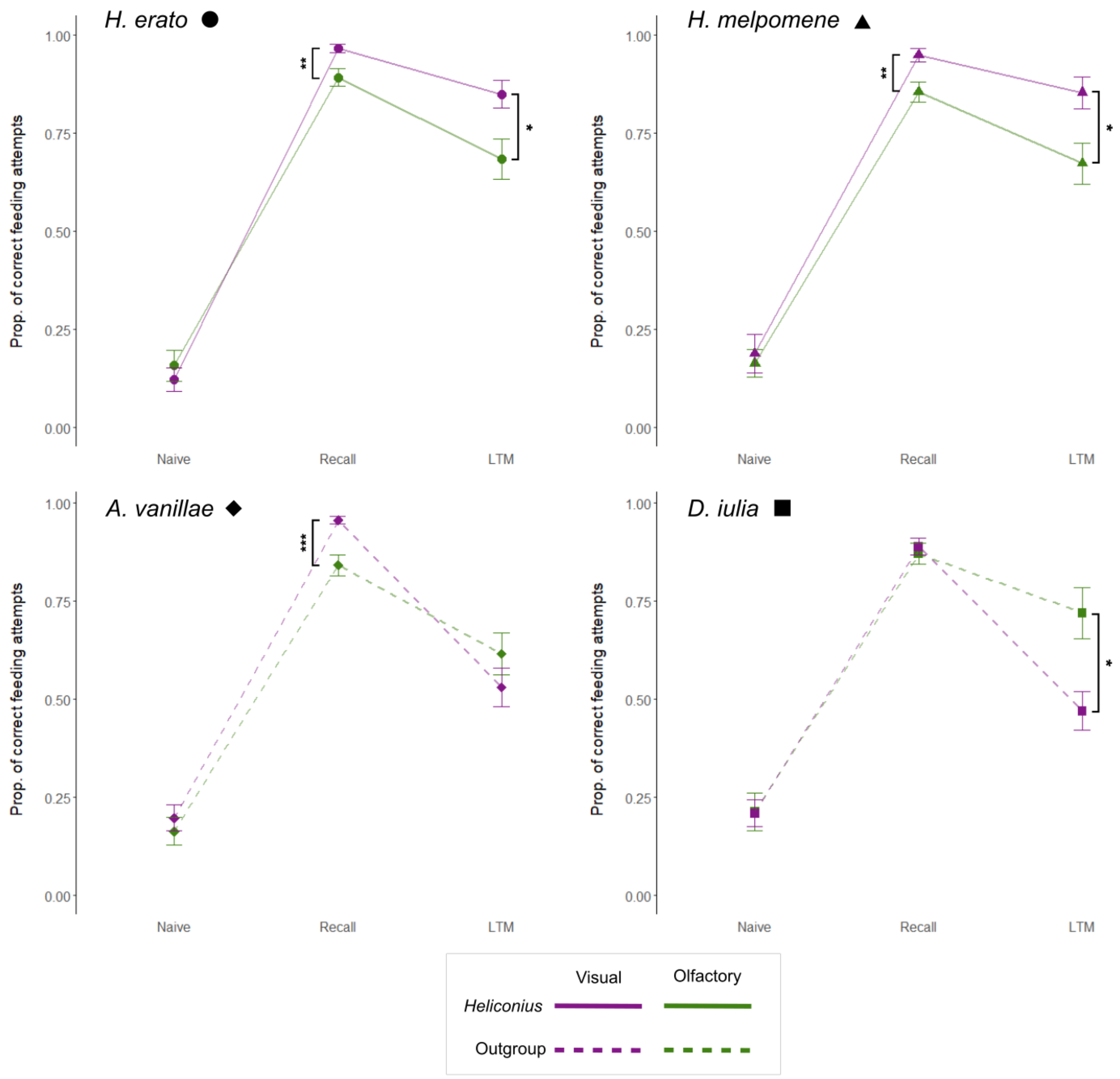


**Figure S5.** Differences in performance across the three trials (naïve, recall and LTM) in the visual (magenta) and olfactory (green) experiments. Values, including standard errors, extracted from coefficients of a GLMM including a significant *Species***Trial***Experiment* interaction. Asterisks indicate significant pairwise differences in performance between the visual and olfactory experiments. *Heliconius* species are shown with solid lines and outgroup Heliconiini species with dashed lines.

**Table S8.** Pairwise comparisons of trial effects on cue preference in the conflict assay for the three different species: *H. erato*, *H. melpomene*, and *D. iulia.*

| Species | Contrast | estimate | z. ratio | p. value |
|---|---|---|---|---|
| *D. iulia* | Recall - Conflict | 4.892 | 6.494 | **<.001** |
| | Recall - Colour | 4.125 | 4.687 | **<.001** |
| | Conflict - Colour | -0.766 | -0.922 | 0.626 |
| *H. erato* | Recall - Conflict | 4.338 | 8.116 | **<.001** |
| | Recall - Colour | 3.004 | 4.900 | **<.001** |
| | Conflict - Colour | -1.334 | -2.435 | **0.040** |
| *H. melpomene* | Recall - Conflict | 5.230 | 5.486 | **<.001** |
| | Recall - Colour | 3.072 | 2.907 | **0.010** |
| | Conflict - Colour | -2.157 | -2.440 | **0.039** |

**Table S9.** Pairwise comparisons between the three species *H. erato*, *H. melpomene*, and *D. iulia*, on cue preference for the three main tests in the conflict assay.

| Trial | Contrast | estimate | z. ratio | p. value |
|---|---|---|---|---|
| Recall | *D. iulia – H. erato* | -0.440 | -0.654 | 0.999 |
| | *D. iulia – H. melpomene* | -1.271 | -1.338 | 0.920 |
| | *H. erato – H. melpomene* | -0.831 | -0.938 | 0.991 |
| Conflict | *D. iulia – H. erato* | -0.995 | -1.676 | 0.762 |
| | *D. iulia – H. melpomene* | -0.933 | -1.268 | 0.941 |
| | *H. erato – H. melpomene* | 0.061 | 0.098 | 1 |
| Colour | *D. iulia – H. erato* | -1.562 | -1.891 | 0.620 |
| | *D. iulia – H. melpomene* | -2.324 | -2.334 | 0.322 |
| | *H. erato – H. melpomene* | -0.762 | -0.900 | 0.993 |

**Table S10.** Proportion of feeding attempts in each trial of the conflict assay made on the trained colour that are above or below 50% for each species, *H. erato*, *H. melpomene*, and *D. iulia*.

| Species | Trial | estimate | Pr(>\|z\|) |
|---|---|---|---|
| *D. iulia* | Recall | 7.737 | **0.001** |
| | Conflict | -0.262 | 0.765 |
| | Colour | 0.751 | **0.007** |
| *H. erato* | Recall | 5.289 | **<.001** |
| | Conflict | 0.927 | **<.001** |
| | Colour | 2.323 | **<.001** |
| *H. melpomene* | Recall | 6.595 | **0.006** |
| | Conflict | 0.982 | 0.126 |
| | Colour | 3.185 | **<.001** |

**Table S11**. Pairwise comparisons between different species, *D. iulia, H. erato* and *H. melpomene*, of the number of feeding attempts made per individual per day in the conflict assay.

| Trial | Contrast | estimate | z. ratio | p. value |
|---|---|---|---|---|
| Naïve | *D. iulia – H. erato* | -0.396 | -1.632 | 0.898 |
| | *D. iulia – H. melpomene* | -0.297 | -0.979 | 0.998 |
| | *H. erato – H. melpomene* | 0.100 | 0.392 | 1 |
| Recall | *D. iulia – H. erato* | -0.609 | -3.273 | **0.049** |
| | *D. iulia – H. melpomene* | -0.695 | -2.907 | 0.138 |
| | *H. erato – H. melpomene* | -0.086 | -0.390 | 1 |
| Conflict | *D. iulia – H. erato* | -1.280 | -5.627 | **<.001** |
| | *D. iulia – H. melpomene* | -1.211 | -4.332 | **0.001** |
| | *H. erato – H. melpomene* | 0.070 | 0.291 | 1 |
| Colour | *D. iulia – H. erato* | -0.557 | -1.829 | 0.802 |
| | *D. iulia – H. melpomene* | -0.306 | 0.875 | 0.999 |
| | *H. erato – H. melpomene* | 0.251 | 0.881 | 0.999 |

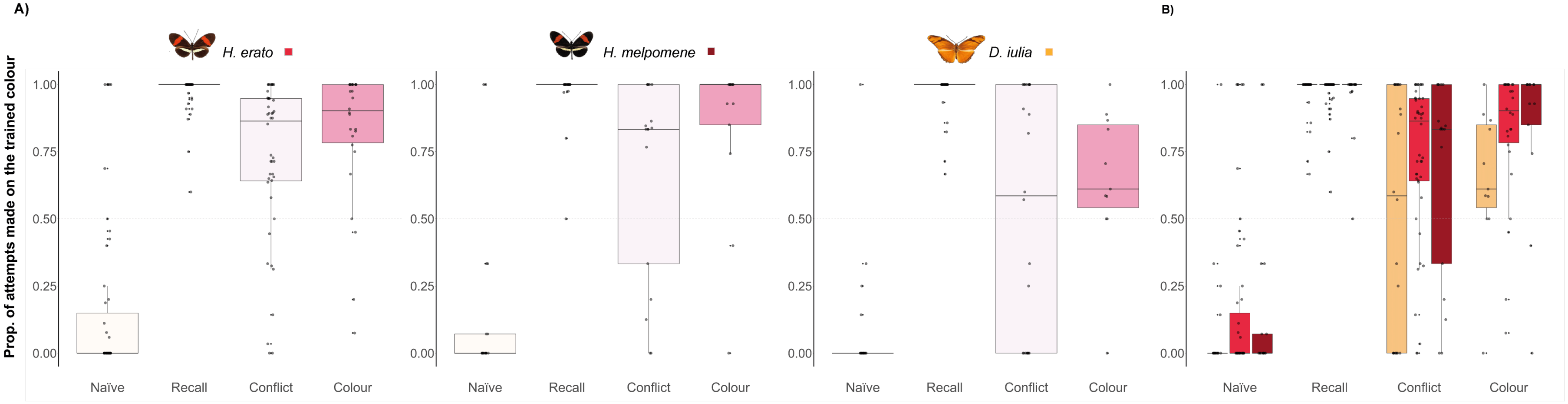


**Figure S6.** Proportion of feeding attempts made on the trained colour by *H. erato*, *H. melpomene*, and *D. iulia* across the different tests, excluding the attempts made on the odour-well Eppendorf tubes. The graphs are colour-coded by test: white for naïve preference, dark orange for the recall test, light pink for the conflict test, and pink for the colour test. In the last graph is colour-coded by species, orange for *D. iulia*, light red for *H. erato* and red for *H. melpomene*. (A) Shows the performance of each species across the different tests. (B) Compares the performance of the three species across the four days.

**Table S12.** Pairwise comparisons of trial effects on cue preference for the three different species: *H. erato*, *H. melpomene*, and *D. iulia*. Excludes the feeding attempts made directly on the odour containing Eppendorf tubes.

| Species | Contrast | estimate | z. ratio | p. value |
|---|---|---|---|---|
| *D.iulia* | Recall - Conflict | 4.625 | 5.958 | **<.001** |
| | Recall - Colour | 3.818 | 4.258 | **<.001** |
| | Conflict - Colour | -0.807 | -0.960 | 0.602 |
| *H. erato* | Recall - Conflict | 4.223 | 6.825 | **<.001** |
| | Recall - Colour | 2.847 | 4.071 | **<.001** |
| | Conflict - Colour | -1.376 | -2.236 | 0.065 |
| *H.melpomene* | Recall - Conflict | 4.446 | 5.180 | **<.001** |
| | Recall - Colour | 2.490 | 2.491 | **0.034** |
| | Conflict - Colour | -1.956 | -2.231 | 0.066 |

**Table S13.** Pairwise comparisons between the three species, *H. erato*, *H. melpomene*, and *D. iulia*, on cue preference for the three main tests. Excludes the feeding attempts made directly on the odour containing odour-well Eppendorf tubes.

| Trial | Contrast | estimate | z.ratio | p. value |
|---|---|---|---|---|
| Recall | *D. iulia – H. erato* | -1.035 | -1.362 | 0.912 |
| | *D. iulia – H. melpomene* | -0.809 | -0.885 | 0.994 |
| | *H. erato – H. melpomene* | 0.226 | 0.256 | 1 |
| Conflict | *D. iulia – H. erato* | -1.438 | -2.120 | 0.460 |
| | *D. iulia – H. melpomene* | -0.988 | -1.262 | 0.942 |
| | *H. erato – H. melpomene* | 0.449 | 0.638 | 0.999 |
| Colour | *D. iulia – H. erato* | -0.631 | -0.740 | 0.351 |
| | *D. iulia – H. melpomene* | -2.137 | -0.198 | 1 |
| | *H. erato – H. melpomene* | -0.131 | -0.145 | 1 |
